# Behavioural individuality determines infection risk in clonal ant colonies

**DOI:** 10.1101/2023.01.26.525341

**Authors:** Z. Li, E.T. Frank, T. Oliveira-Honorato, F. Azuma, V. Bachmann, D. J. Parker, T. Schmitt, E. Economo, Y. Ulrich

## Abstract

In social groups, disease risk is not distributed evenly across group members. Individual behaviour is a key source of variation in infection risk, yet its effects are difficult to separate from those of other factors. Here, we combine long-term epidemiological experiments and automated tracking in clonal raider ant colonies, where behavioural individuality emerges among identical workers. We find that: 1) division of labour determines the distribution of parasitic nematodes (*Diploscapter*) among hosts, showing that differences in infection can emerge from behavioural variation alone, 2) infections affect colony social organisation by causing infected workers to stay in the nest. By disproportionally infecting some workers and shifting their spatial distribution, infections reduce division of labour and increase spatial overlap between hosts, which is expected to facilitate parasite transmission. Thus, division of labour, a defining feature of many societies, not only shapes infection risk and distribution but can also be modified by parasites.

## Introduction

Host spatial behaviour is a key driver of parasite infection and transmission success. For example, host spatial distribution modulates infection risk, with feeding, breeding, or resting sites often acting as disease transmission hotspots^1,2^. Infections can in turn have drastic effects on host spatial behaviour, ranging from manipulations that increase parasite transmission to new hosts to host behavioural responses that reduce transmission. For example, some pathogenic fungi are thought to increase their transmission by causing infected ants to climb up and bite vegetation before dying so fungal spores are released from their cadaver onto foraging trails after their death^3^. Conversely, human and insect societies alike contain the spread of pathogens by reducing spatial overlap between group members^4,5^. Empirical studies on the relationship between host spatial behaviour and parasite infection risk largely rely on observations of natural populations^6–8^, which are instrumental in generating hypotheses but can be confounded with other factors^9,10^. Experimental tests of the link between host spatial behaviour and infection risk, however, are comparatively rare, in part because few systems allow controlled infection experiments and monitoring of individual behaviour in replicate host groups of known composition.

Social insect colonies are powerful systems to study the link between behaviour and infection because they display complex yet tractable spatial behaviour. While the core of the colony (the nest) is usually static, colony members exhibit heterogeneous spatial distribution around the nest that reflects their behavioural roles: for example, nurses mainly stay in the nest to care for the brood, while foragers leave the nest to collect food^11,12^. This spatial organisation of division of labour^11^ has in turn been proposed to result in asymmetries in infection risk, with workers specialising in tasks outside the nest (e.g., foraging, waste management) expected to have increased exposure to parasites present in the external environment. While this assumption is central to the study of disease transmission in social insect colonies^13,14^, it is difficult to test rigorously. This is in part because behavioural roles in most social insect colonies covary with several individual traits, including age^11^ and genotype^15^. These factors in turn typically affect disease resistance independently from behaviour^16–18^, making it difficult to unambiguously measure the effects of behaviour on infection risk.

To overcome this problem, we investigate the link between behaviour and parasitic infection in the clonal raider ant (*Ooceraea biroi*). Colonies of this species are queenless and composed of genetically near-identical workers that reproduce asexually and synchronously^19^, providing experimental control over genotype and age at the individual and group levels. Furthermore, colonies composed of identical individuals show spatially organised division of labour^20^, i.e. individuality in spatial behaviour that reflects behavioural roles, making it possible to study the link between host behaviour and parasite infection in the absence of confounding variation in age or genotype.

As an infectious agent, we use a nematode of the genus *Diploscapter*—the sister group of *Caenorhabditis*—that we find naturally infects *O. biroi* and other ants. Nematodes of this and related genera have sporadically been reported to infect ants since the late 19^th^ century^21–26^, but the nature of their interaction with ants remains poorly characterized. Strikingly, the association between nematodes and ants displays high organ-specificity: nematodes were repeatedly reported to infect a specific head gland, the pharyngeal gland (PG)^21–25^. The PG, previously called the postpharyngeal gland until recent reassessment^27^, has an eminently social function in ants, with roles in the transfer of nutrients and signalling molecules between colony members^28,29^. In particular, the PG plays a key role in the exchange of cuticular hydrocarbons (CHC) across colony members^30^. In ants, CHC profiles form the chemical basis for nestmate vs. non-nestmate discrimination, and are associated with behavioural roles^31^. Thus, while *Diploscapter* nematodes have often been assumed to use ants as a mere means of dispersal^22,24,25^, the localisation of nematodes in the PG suggests that infections might affect the social organisation of the colony.

## Results

### *Diploscapter* naturally infects ants and affects host fitness and physiology

Sequencing confirmed that nematodes isolated from the heads of workers from *O. biroi* and two other ant species (*Paratrechina longicornis* and *Lasius niger*, Fig. 1) cluster with *Diploscapter* sampled from the body and nest material of *Prolasius advenus* ants in New Zealand^26^ (Fig. 1a). Thus, together with previous reports^22,24,25^, our findings support the view that some *Diploscapter* nematodes naturally infect ants^32^. Micro-CT reconstructions of infected ant heads confirmed that *Diploscapter* localises to the PG in the studied species (Fig. 1b). Although the PG varies in shape and arrangement across species, in each case the nematodes fill the lumen of the gland, where they can reach high densities and become tightly packed in bundles within the different “fingers” of the glove-shaped PG.

**Figure 1.**
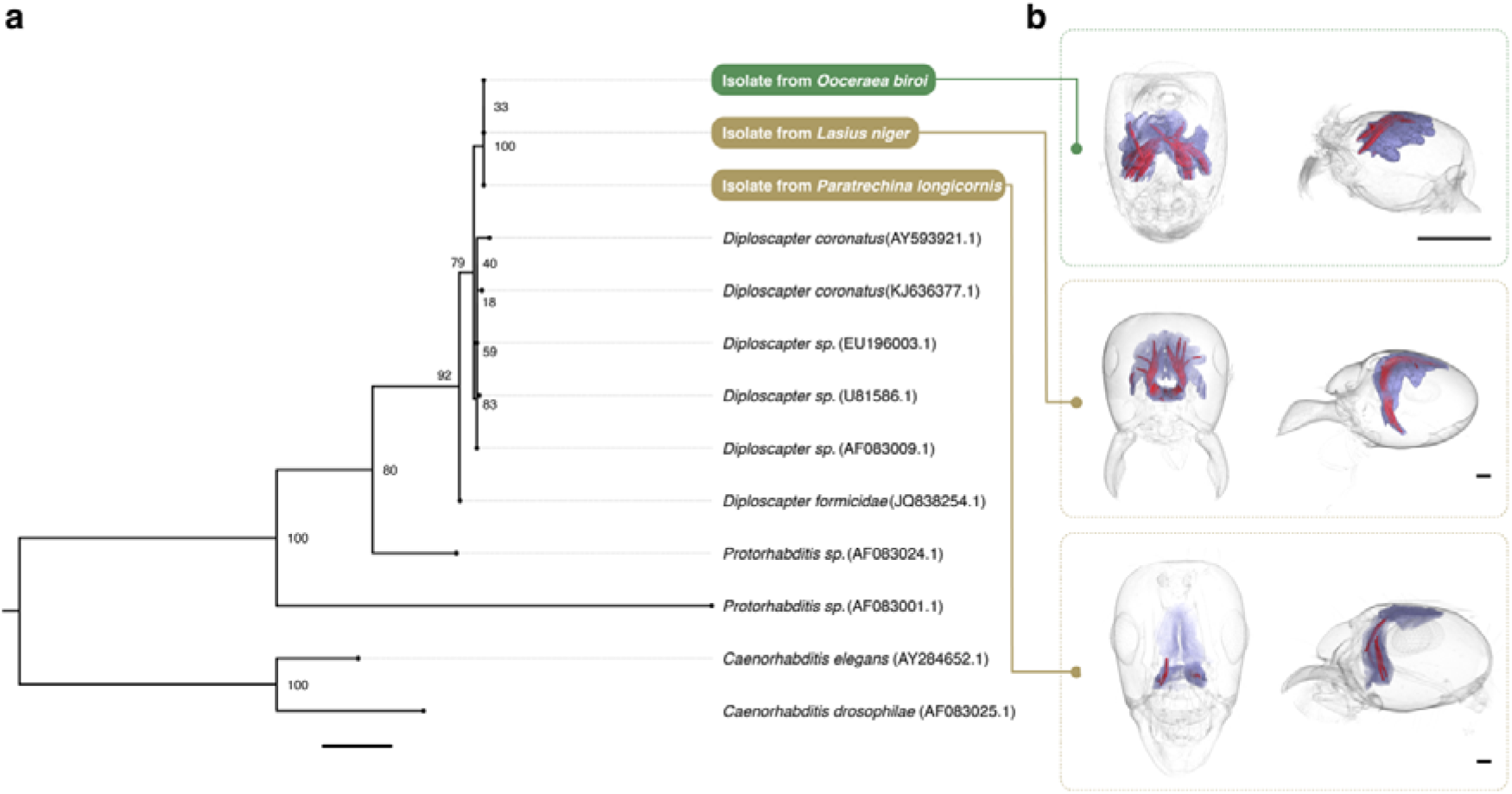
*Diploscapter* nematodes infect the PG of ants. (a) *Diploscapter* isolated from the heads of clonal raider ants (green) and two other ant species (yellow) are phylogenetically nested within the *Diploscapter* clade (sequence accession numbers in brackets). Node labels indicate branch support (%) from 1000 bootstrap replicates. Scale bar: mean number of 0.02 substitutions per site. (b) Micro-CT reconstructions show that nematodes (red) localise to the PG (purple) in all studied ant species. Scale bars: 0.2 mm.

While previous studies proposed a phoretic relationship between *Diploscapter* nematodes and ants^22,24,25^, our results indicate that these nematodes do not use their ant hosts for dispersal alone. First, infections affected host survival. Experimentally infected clonal raider ant workers had reduced survival (mean ± SD: 75.00 ± 4.84%) relative to otherwise identical uninfected workers (96.88 ± 1.78%) over 51 days (Cox proportional-hazards model: hazard ratio = 9.96, z = 3.40, p = 0.0007; Fig. 2a). Second, infections strongly affected gene expression in the PG, including immune genes. We detected 657 differentially expressed genes in the PG (Tab. S1) of clonal raider ants but no differentially expressed genes in an adjacent uninfected organ, the brain (Fig. 2b, Tab. S2). Gene set enrichment analyses showed enrichment for a diverse set of gene ontology (GO) terms (n = 333), showing infection affected a broad range of functional processes. To aid interpretation we semantically clustered enriched GO terms and found 8 clusters with clear links to immune responses (e.g., defence response, response to bacterium, wound healing), as well as 9 clusters with clear links to behavioural processes (e.g., regulation of behaviour, social behaviour, aggressive behaviour) (Tab. S3). The majority of DE genes in immune-related clusters showed increased expression in infected individuals (Fig. S1, Tab. S3), consistent with an upregulation of the immune response. This effect was largest for genes related to wound healing (9/11 genes were upregulated) suggesting infection may cause physical damage to the PG. Lastly, infections affected host CHC profiles (ADONIS: R^2^= 0.62, F=76.79, p <0.0001; Fig. 2c, Tab. S4). Our chemical analysis showed that infections altered the relative abundance of all CHC classes on the cuticle of clonal raider ant workers: both *n-*alkanes and methyl-branched alkanes had lower relative abundance in infected individuals (Mann–Whitney U tests: *n*-alkanes: U = 5, p <0.0001; methyl-branched alkanes: U = 74, p <0.001), while dimethyl-branched alkanes had greater relative abundance (U = 382, p <0.0001) (Fig. 2d). Together, our findings show that some nematodes of the genus *Diploscapter* are facultative parasites of ants that affect the physiology and reduce the fitness of their hosts.

**Figure 2.**
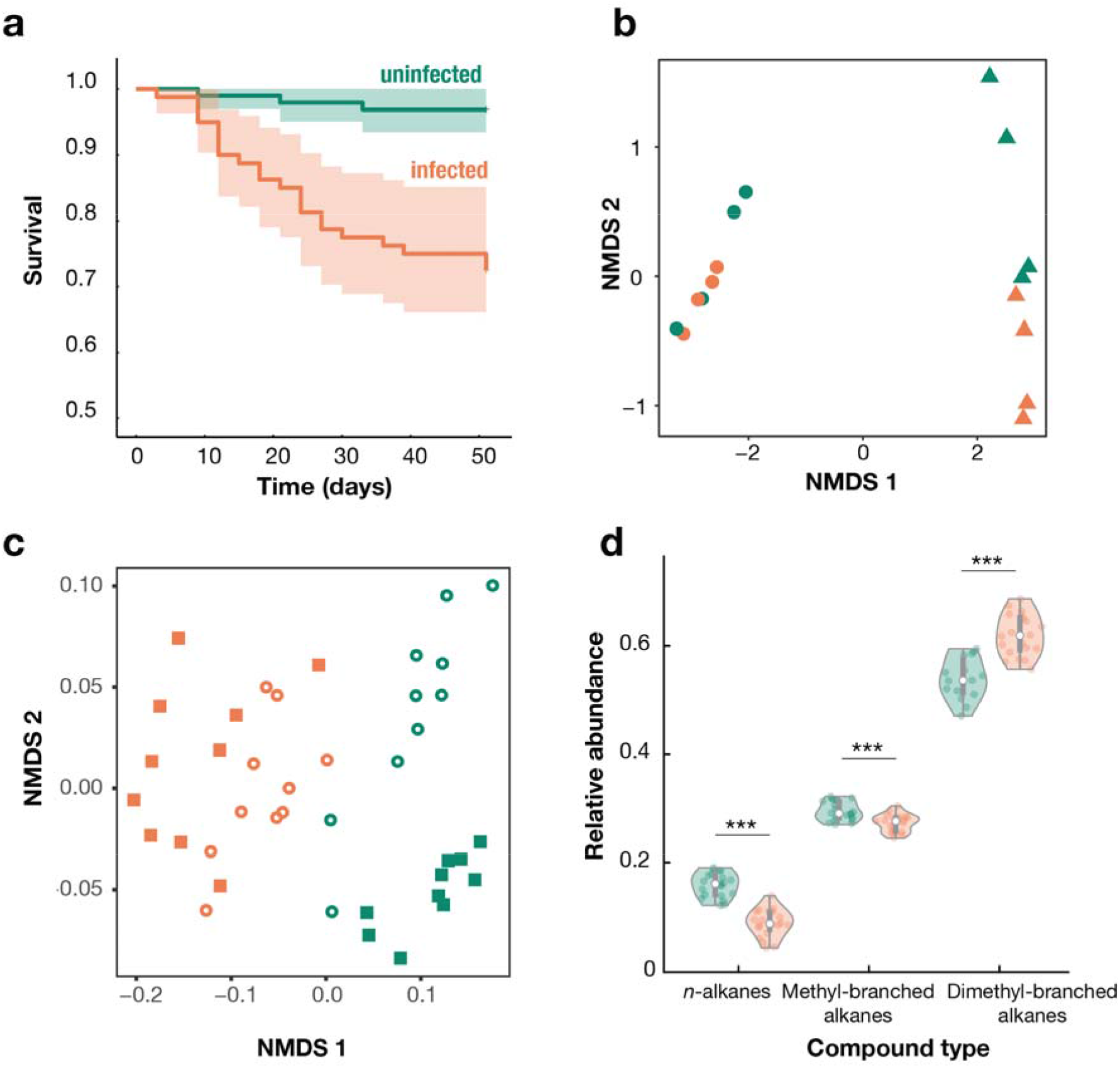
Nematode infections affect host survival and physiology. (a) Survival of infected (n = 80 workers in 10 colonies) and uninfected (n = 96 workers in 12 colonies) ants over time. Lines and shaded areas represent means and 95% confidence intervals. (b) Gene expression in the brain (circles) and PG (triangles) of infected (orange) and uninfected (green) ants. Data points represent pooled samples of 5 ants. (c) CHC profiles of infected (orange) and uninfected (green) ants from genetic lineages B (open circles) and L (squares). Data points represent pooled samples of 5 decapitated ants. (d) Relative abundance of CHC classes in infected (orange) and uninfected (green) ants. Thick grey bars indicate the interquartile range around the median, and thin grey bars represent 1.5 times the interquartile range. Data points represent pooled samples of 5 decapitated ants. ***: p <0.001.

### Behavioural individuality determines infection risk

Like other nematodes^33,34^, *Diploscapter* finds hosts by nictation^33^, a stereotypical waving behaviour that allows attachment to passing hosts. We hypothesized that hosts that differ in spatial behaviour (but are otherwise identical) may be at different risk of infection with such ambush-style parasites. We tested this hypothesis by exposing uninfected *O. biroi* colonies made up of age-matched, clonally-related workers to agar seeded with nematodes over four colony cycles (216 days). Nictating nematodes attached to ants walking within their reach (Mov. S1). As predicted, we find that infection risk is not homogenously distributed across colony members, but instead depends on individual behavioural roles. During the first colony cycle, nematode prevalence was already higher (mean ± SD: 90.00 ± 30.5%) in the workers acting as foragers outside the nest than in the otherwise identical workers acting as nurses in the nest (33.33 ± 47.9%) (generalised linear mixed model (GLMM): worker behavioural role, degrees of freedom (DF) = 1, likelihood ratio (LR) = 22.19, p <0.0001; Fig. 3a). In the subsequent cycles, all sampled ants were infected, but infection load increased faster in foragers than in nurses (GLMM: interaction between time and worker behavioural role, DF = 2, LR = 51.42, p <0.0001; Fig. 3a). While in the second cycle, the mean infection loads of foragers (12.80 ± 10.80 nematodes/ant) and nurses (11.60 ± 9.09) were undistinguishable (pairwise comparison of estimated marginal means (EMM): z = 1.33, p = 0.18), in the third and fourth cycles, foragers had drastically higher infection loads than nurses (third cycle: foragers: 70.70 ± 32.20, nurses: 35.30 ± 17.40; z = 18.44, p <0.0001; fourth cycle: foragers: 86.40 ± 31.00, nurses: 46.90 ± 23.00; z = 18.32, p <0.0001). Thus, the distribution of parasites across hosts reflected inter-individual variation in host spatial behaviour driven by age- and genotype-independent division of labour, showing that differences in infection can emerge from differences in behaviour alone.

**Figure 3.**
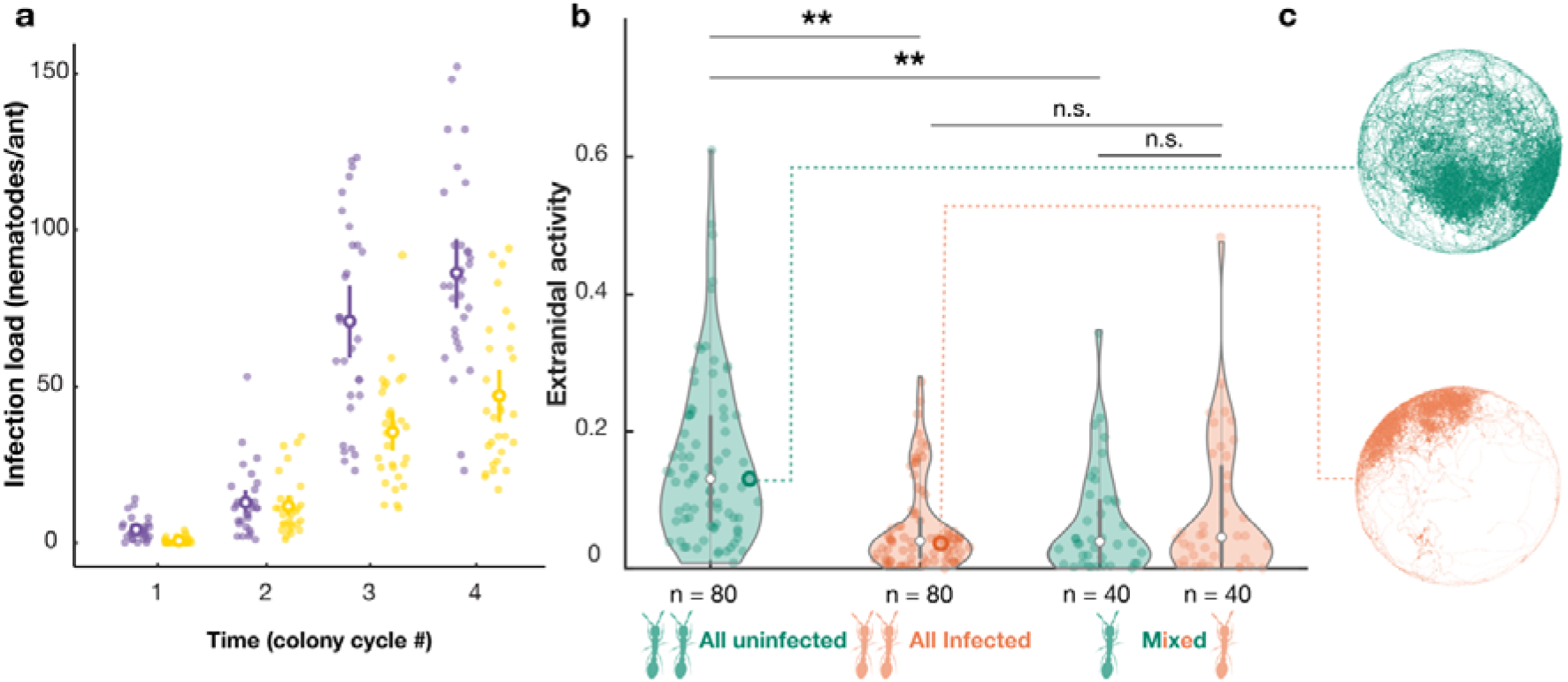
Effects of behaviour on infection, and vice versa. (a) Infection load in foragers (purple) and nurses (yellow) over time. Small opaque dots represent individual workers (n = 30 per behavioural role and colony cycle). Large circles and bars represent mean ± s.e. (b) Extranidal activity (proportion of time outside the nest) of uninfected (green) and infected (orange) workers as a function of colony infection composition. Thick grey bars indicate the interquartile range around the median, and thin grey bars represent 1.5 times the interquartile range. Opaque dots and sample sizes represent individual ants. Shaded areas are kernel density estimates. (c) Spatial distribution of representative workers (open circles in b) from an uninfected and an infected colony. **: p <0.01, n.s.: not significant.

### Infections alter social organisation

Using automated tracking of experimentally infected hosts over 6 days, we find that not only does host spatial behaviour affect infection risk (Fig. 3a), but infection also affects host spatial behaviour (GLMM, DF = 3, LR = 11.24, p = 0.01; Fig. 3bc). Workers in infected colonies spent more time in the nest (i.e., behaved more like nurses) than otherwise identical workers in uninfected colonies (extranidal activity: Infected_all_: EMM (95% confidence interval): 0.06 (0.04, 0.09), Uninfected_all_: 0.14 (0.10, 0.20); t = 3.14, p = 0.008). Unexpectedly, the presence of some infected individuals in the colony affected the behaviour of all colony members: the extranidal activity of uninfected workers in mixed colonies was as low as that of their infected nestmates (Infected_mixed_: 0.06 (0.04, 0.10), Uninfected_mixed_: 0.06 (0.04, 0.09), t = 0.46, p = 0.983)—and lower than that of uninfected ants in nematode-free colonies (Uninfected_all_ vs. Uninfected_mixed_, t = 3.14, p = 0.008)—although they remained uninfected throughout the experiment (i.e., no transmission occurred, as confirmed by dissections). In contrast, the extranidal activity of infected ants was not increased by the presence of uninfected ants (Infected_all_ vs. Infected_mixed_: t = -0.14, p = 0.999). Because nematodes disproportionally infect foragers (Fig. 3a) and infections induce nurse-like spatial behaviour (Fig. 3b), our results imply that at the colony-level, exposure to nematodes should effectively dampen division of labour.

## Discussion

We report *Diploscapter* nematodes in the PG of three ant species and show that infections affect host fitness, physiology, and behaviour in *O. biroi*. Our findings support the view that some *Diploscapter* nematodes represent an early, incomplete transition to parasitism^32^ in a clade of otherwise free-living, bacterivore nematodes related to *C. elegans*^35^. The association of *Diploscapter* with ants is geographically widespread and organ-specific, but the nematodes retain the ability to complete their life cycle without hosts^25^. However, infections induce an immune reaction and decrease host survival, indicating that these nematodes do not merely use ants for dispersal, as previously thought^22,24,25^, but may instead derive resources from their host. We suggest that the contents of the PG make it an attractive source of nutrients for nematodes. The shift in host CHC profiles in infected workers, including reduced ratios of *n*-alkanes and methyl-branched alkanes, suggests nematodes may use hydrocarbons as a carbon source, potentially with a preference for certain compounds. In addition, the PG of ants contains sterols^29,36^, which *C. elegans* requires exogenously for survival and to develop from dauer larvae (the persistent larval stage that is also the infective stage in many parasitic nematodes) into reproductive adults^37^. Further work is needed to establish what resources (if any) *Diploscapter* nematodes derive from ant PGs, and what role they play in their development.

Combining semi-natural and experimental infections with behavioural analyses, we show that *Diploscapter* nematodes interact with a defining feature of insect societies, division of labour. Workers active outside the nest became infected earlier and carried higher infection loads than otherwise identical individuals acting as nurses in the nest. Thus, differences in infection can result from differences in behaviour alone and the uneven distribution of parasites across group members can emerge from the spatial organization of division of labour in the colony. This in turn highlights colony behavioural composition as a key driver of infection dynamics. Previous work in the clonal raider ant has shown that colony size^20^, as well as colony demographic, genetic, and morphological composition^38^ affect the spatial organisation of division of labour. Our results predict that each of these aspects of colony composition should in turn affect the dynamics of parasite acquisition in the colony. The optimal allocation of individuals to tasks in social insect colonies might therefore reflect a trade-off between the benefits (e.g., nutrient intake) and costs (e.g., parasite acquisition) of extranidal activity.

Not only did host spatial behaviour affect infection probability, but infections in turn affected host spatial behaviour at both the individual and colony levels. Experimentally infected workers spent considerably more time in the nest in proximity to the brood and nestmates. Surprisingly, the presence of infected ants also reduced the extranidal activity of their uninfected nestmates, potentially through social processes. Clonal raider ants forage in scout-initiated group raids, with one or a few scouts recruiting their nestmates to food outside the nest^39^. An infection-induced reduction of extranidal activity in some colony members would reduce the foraging activity of other colony members if it decreases the occurrence of scout-initiated raids. The observed reduction of extranidal activity upon infection contrasts with behavioural effects of infection with well-studied fungal pathogens, where infected hosts tend to reduce interactions with nestmates^4^ or to leave the nest altogether^40–42^. Here, the infection-induced changes in host behaviour increased spatial overlap between hosts and are therefore expected to increase the nematode’s transmission success. From the parasite’s perspective, a host that spends time in the nest presents more transmission opportunities than a forager (irrespective of whether transmission occurs via direct contact or through the environment) because the nest is invariably the place where host density is highest.

Taken together, our behavioural analyses show that extranidal workers are more likely to become infected but that infections in turn reduce extranidal activity. Thus, by turning extranidal workers into intranidal workers, nematode infections effectively dampen division of labour at the colony level. Division of labour is thought to provide an organizational barrier against disease transmission in social insects because it creates spatial segregation between behavioural groups that can compartmentalize infections and slow disease spread^43,13,44,45^. By disrupting division of labour and increasing spatial overlap between colony members, infecting nematodes may collapse that organisational barrier to their own advantage.

## Materials and Methods

### Ants and Nematodes

Colonies of the clonal raider ant are queenless and consist of genetically near-identical, totipotent workers that reproduce asexually and synchronously, producing new workers in discrete cohorts^19,46^. This provides the ability to create replicate experimental colonies with precise control of genetic and demographic variation both within and between colonies. Synchronized reproduction drives stereotypical colony cycles lasting ca. 5 weeks^47^, in which colonies alternate between reproductive and brood-care phases, corresponding to the absence and presence of larvae, respectively. During the reproductive phase, all ants remain in the nest and lay eggs. During the brood-care phase, the ants attend to the growing larvae at the nest but also leave the nest to forage, explore, or dispose of waste. Colonies containing as little as 6 ants are fully functional and have division of labour under laboratory conditions in this species^20^.

Nematodes were isolated from workers of laboratory colonies of *O. biroi* originally collected in Okinawa and the US Virgin Islands^48^. Because we cannot rule out transmission between colonies after collection, the exact geographical origin of the *O. biroi* nematode isolate cannot be confirmed. Two further isolates were obtained from workers of *Paratrechina longicornis* collected in Malaga, Spain, and *Lasius niger* workers sampled in Lausanne, Switzerland.

Nematodes were isolated from dissected ant heads of each species and grown on agar plates seeded with bacteria (*Escherichia coli* OP50)^49^. To prepare infection inocula, nematodes were washed from agar plates with ddH_2_O. The concentration of the resulting solution was determined (average of 3 independent counts per sample) in counting chambers (MultiCount10, Immune Systems Ltd., UK) and adjusted to the desired concentration (see below). Because infections are localised to the PG and live *Diploscapter* nematodes are not found elsewhere in the ant body, the infection status of individual ants can reliably be assessed by dissecting their head. Nematodes do not infect the brood, as shown by the fact that uninfected colonies can be established with newly-eclosed callow workers from infected colonies and remain uninfected for years. All experiments were performed with the *Diploscapter* isolate from *O. biroi* ants.

### Nematode phylogenetic analysis

We Sanger sequenced the SSU (18S rDNA) of three nematode isolates following Holterman et al.^50^. Sanger sequences were trimmed with Geneious Prime (v. 2021.0.3) to produce a consensus sequence. We then downloaded all available *Diploscapter* 18S sequences with >1000 bp of sequence from NCBI, as well as two *Protorhabditis* and two *Caenorhabditis* sequences, which are outgroups of *Diploscapter*^50^. Sequences were aligned using PRANK (v.100802, default options)^51^. The alignment was cleaned with Gblocks (v. 0.91, minimum block length = 50)^52^ to eliminate poorly aligned positions. We then used RAxML (v. 8.2.12, options -m GTRGAMMA -p 12345 -# 1000 -f a -x 12345 -c 40 -T 10)^53^ to generate a maximum-likelihood tree, assuming a GTR + gamma model of sequence evolution with 1000 bootstrap iterations.

### Micro-CT scans and segmentation

One worker of the infected *Lasius niger, Ooceraea biroi*, and *Paratrechina longicornis* were scanned and each given a unique specimen identifier (Tab. S5).

The specimens were fixed in 70% ethanol, stained in 2M iodine solution for a minimum of 7 days, and then washed and individually sealed in a pipette tip with 99% ethanol for scanning. The scans were performed with the ZEISS Xradia 510 Versa 3D X-ray microscope at the Okinawa Institute of Science and Technology, Japan.

The head of each specimen was scanned at 40kV and 3W for a full 360º rotation with 1601 projections (except *O. biroi* which had 3201 projections). Exposure time was set to yield intensity levels between 13,000-15,000 across the head and ranged from 13-24.5 seconds. The resulting scans were reconstructed with the ZEISS Scout-and-Scan Control System Reconstructor software (ZEISS, Oberkochen, Germany).

The PG and parasites within the gland were manually segmented using the brush tool in Amira (v2019.2). The segmented materials were exported as Tiff image stacks, imported into VGStudio (v3.4; Volume Graphics GmbH, Heidelberg, Germany) and visualized using volume rendering (Phong).

### Survival assay

Uninfected, age-matched (77-days old), clonally related (genotype A) workers were divided into two groups: infected (140 ants) and uninfected (140 ants). Each group was kept in a plastic nest box (110 × 150 × 80 mm) with a plaster of Paris floor. Ants from the uninfected group were kept nematode-free, while ants from the infected group were exposed to nematodes by the addition of 10-100 infected ant heads in their nest box every 4-11 days. 75 days post-exposure, the success of the treatment was confirmed by dissecting 10 ants per treatment: all ants from the uninfected group were nematode-free and all ants from the infected group carried nematodes (6.8 ± 1.29 nematodes/ant). Ants were split into 10 infected and 12 uninfected experimental colonies in airtight Petri dishes (diameter 50mm) with a plaster of Paris floor. Each experimental colony consisted of 8 ants and 6 larvae. Every 3 days over 51 days, colonies were watered and fed with frozen ant (*Messor*) brood, worker survival was monitored, and any dead ants were replaced with genotype- and age-matched ants from the corresponding treatment (uninfected or infected) to rule out effects of declining group size on survival.

### Transcriptomic analysis

Brains and PGs were dissected from 20 infected and 20 uninfected age-matched (85-days old), clonally related (genotype A) workers. The workers had been exposed to nematodes on agar (infected) or nematode-free agar (uninfected) as a group for 55 days prior to dissection. Brains or PGs of 5 workers were pooled to produce 4 samples per tissue type. RNA was extracted from each sample using a modified Trizol/phenol chloroform protocol following Libbrecht et al.^54^. Libraries were prepared using a KAPA stranded mRNA-sequencing kit following the manufacturer’s protocol. Libraries were sequenced using standard protocols on an Illumina HiSeq 4000. Raw sequencing reads have been deposited in NCBI’s sequence read archive under the following BioProject accession: PRJNA791185.

Reads were trimmed using Trimmomatic (v. 0.36)^55^, to remove adaptors and low-quality bases (LEADING:9 TRAILING:9 SLIDINGWINDOW:4:15). Reads less than 90bp after trimming were discarded. Trimmed reads were then mapped to transcripts extracted from the *O. biroi* reference genome (GCF_003672135.1_Obir_v5.4^56^) with Kallisto (v. 0.46.0)^57^. Read counts for each gene were then obtained using tximport (v. 1.10.1) ^58^ in R (v4.0.3) (Tab. S6). Expression analyses were performed using the Bioconductor package EdgeR (v. 3.24.0)^59^. rRNA genes and genes with counts per million < 0.5 in two or more libraries per condition were excluded. Expression analyses were performed separately for each tissue. Normalization factors for each library were computed using the TMM method. To estimate dispersion, we fit a generalized linear model with a negative binomial distribution with infection status as an explanatory variable. A quasi-F test was used to determine the significance of infection status from this model for each gene, with p values corrected for multiple tests using Benjamini and Hochberg’s^60^ algorithm. Statistical significance was set to 5%.

Genes were functionally annotated using Blast2GO^61^ as follows: Gene sequences were compared with blastX to either NCBI’s nr-arthropod or *Drosophila melanogaster* databases, keeping the top 20 hits with e-values <1 × 10^−03^. Interproscan (default settings within Blast2GO) was then run for each sequence, and the results merged with the BLAST results to obtain GO terms. This produced two sets of functional annotations, one derived from all arthropods and one specifically from *D. melanogaster*. The *D. melanogaster* GO term annotation generated around four times more annotations than NCBI’s nr-arthropod database. We therefore conducted all subsequent analyses using the GO terms derived from *D. melanogaster* but note that enrichment analyses using the annotations from all arthropods found broadly similar terms albeit with fewer and less specific annotations. Overrepresented GO terms were identified by conducting gene set enrichment analyses (GSEA) using the R package TopGO (v. 2.28.0)^62^, using the elim algorithm to account for the GO topology. GO terms were considered to be significantly enriched when p <0.05. Enriched GO terms were then semantically clustered using ReviGO^63^ to aid interpretation.

### Chemical analysis

To quantify differences in CHC profiles between infected and uninfected ants, CHCs were sampled from infected and uninfected colonies of two genotypes (B and L; 50 workers for each genotype and infection status combination). All ants were sampled from colonies in the reproductive phase to rule out potentially confounding effects of colony phase on CHC profiles. Heads were dissected to confirm infection status and headless bodies were pooled in groups of 5, producing 10 samples per genotype-infection status combination for GC-MS analyses.

CHCs were extracted from pooled samples by immersion in 1 mL of hexane for 10 min. Extracts were then evaporated to a volume of approximately 15 μL of which 1 μL was analyzed using a 6890 gas chromatograph (GC) coupled to a 5975 mass selective detector (MS) (Agilent Technologies, Waldbronn, Germany). The GC was equipped with a DB-5 capillary column (0.25 mm ID × 30 m; film thickness 0.25 μm, J & W Scientific, Folsom, CA, USA). Helium was used as a carrier gas with a constant flow of 1 mL/min. A temperature program from 60 °C to 300 °C with 5 °C/min and finally 10 min at 300 °C was employed. Mass spectra were recorded in the EI mode with an ionization voltage of 70 eV and a source temperature of 230 °C. The software ChemStation v. F.01.03.2357 (Agilent Technologies) for Windows was used for data analysis. Identification of the compounds was accomplished by comparison of library data (NIST 17) with mass spectral data of commercially purchased standards for *n*-alkanes, diagnostic ions and retention indices. To calculate the relative abundance of CHC compounds, we used the GC abundance of each compound. The area under each peak was quantified through integration and divided against the total area under all CHC-relevant peaks.

### Effects of behaviour on infection

Uninfected, age-matched (74-days old), clonally related (genotype A) ants were divided into three infected colonies (120 ants each) and one control colony (50 ants). The control colony was used to verify that the ants remained nematode-free during the whole period of the study and was not included in statistical analyses. All groups were kept in nest boxes (110 × 150 × 80 mm) with a plaster of Paris floor. For each infected colony, 1.5g of agar was placed in the box and inoculated with 18’960 nematodes in 100 µL water. For the control group, nematode-free agar (1.5g) was used.

In each colony, ants were dissected at 4 time points corresponding to the brood care phase of 4 successive colony cycles. The first time point was 35 days after the start of the experiment for all colonies. After this, cycles became slightly desynchronized across colonies and for each infected colony, the subsequent dissection occurred 8 days after the eclosion of new workers, corresponding to the middle of the brood care phase, when foraging activity is at its peak. At each time point, the heads of 10 workers collected in the nest (“nurses”) and 10 workers collected outside the nest (“foragers”) were dissected under a stereomicroscope and infection load was determined by counting nematodes. Colony size was kept constant by removing all newly eclosed callows at each colony cycle, except for 20 callows that were paint-marked (to be distinguished from the original workers) and returned to the colony to replace the 20 dissected ants. Workers from the uninfected control colony were dissected at each time point to ensure that no contamination took place. The experiment lasted 216 days in total.

### Effects of infection on behaviour

Uninfected, age-matched (20-day old), clonally related (genotype A) ants were divided into 2 groups: infected (150 ants) and uninfected (150 ants). Each group was kept in a plastic nest box (110 × 150 × 80 mm) with a plaster of Paris floor. Ants from the uninfected group were kept nematode-free, while ants from the infected group were exposed to 28’000 nematodes in 100 µL water on 1.5g of agar. 45 days post-exposure, ants were split into 30 experimental colonies in airtight Petri dishes (diameter 50mm) with a plaster of Paris floor: 10 uninfected colonies, 10 infected colonies, and 10 mixed colonies (with half infected and half uninfected workers). Each experimental colony consisted of 8 workers (by that time 65-day old) and six 6-day-old larvae. Workers were tagged with unique combinations of paint marks (Uni Paint PX-20 and PX-21) on the thorax and gaster to be individually detected in video analyses.

Behavioural data were acquired from all experimental colonies over the first 6 days of the experiment. During this period, all colonies were in the brood care phase and no ant died in either treatment. 20 minutes of video (10 frames per second) were recorded every 2 hours throughout this period using webcams (Logitech C910). Every 3 days, the colonies were watered and fed live ant brood (*Messor*). For each colony, the experiment ended when all larvae had eclosed into new adults (i.e., when the colony completed a cycle) or after 60 days (if the colony failed to complete a cycle). At the end of the experiment, the heads of all workers in each colony were dissected and nematodes were counted. Individual trajectories were extracted from videos using the software anTraX^64^. Extranidal activity, defined as the fraction of time an ant was outside the nest, was computed using MATLAB v.2022a (MathWorks, Natick, MA, USA) following Jud et al.^47^.

### Statistical analyses

All statistical analyses were conducted in R 4.1.2^65^.

The effect of treatment (infected vs. uninfected) on individual survival was assessed using a Cox proportional-hazards mixed model (function *coxme* of package *coxme*^66^) with colony as a random effect. Model assumptions were verified using the *cox*.*zph* function from the package *survival*.

CHC profiles were visualised with a nonmetric multidimensional scaling (NMDS) analysis using a Bray-Curtis dissimilarity matrix with the *metaMDS* function from the package *vegan*^67^. The effect of infection status (infected vs. uninfected) on CHC profiles composition was analysed using a permutational multivariate analysis of variance (ADONIS) on a Bray-Curtis dissimilarity matrix with genotype as a random effect with the *adonis* function from the package *vegan*^67^. The relative abundance of different CHC classes (*n*-alkanes, methyl-branched alkanes, dimethyl-branched alkanes) was compared between uninfected and infected ants using Mann-Whitney U tests.

To model nematode prevalence in the first colony cycle, we used a binomial GLMM with logit link function (function *glmer* from package *lme4*) with individual infection status (infected vs. uninfected) as response variable, worker behavioural role (forager vs. nurse) as fixed predictor, and colony as a random effect. To model infection loads in the following cycles, we used a Poisson GLMM with log link function (function *glmer* of package *lme4*) with colony cycle (a three-level factor), worker behavioural role (forager vs. nurse), and their interaction as fixed predictors, and the colony as a random effect. Pairwise comparisons between the estimated marginal mean (EMM) infection loads of foragers and nurses at colony cycles 2 to 4 were performed using posthoc tests implemented with the *emmeans* function from the package *emmeans*^68^. P values were adjusted using the Tukey method for multiple comparisons.

To model worker extranidal activity, a beta GLMM with logit link function was implemented with the function *glmmTMB* from the package *glmmTMB*^69^. The model included a four-level variable combining infection status and colony composition (Uninfected_all_, Infected_all_, Uninfected_mixed_ and Infected_mixed_) as fixed predictor and colony as random effect. Posthoc tests between the EMM extranidal activity between selected groups (Uninfected_all_ vs. Infected_all_, Uninfected_mixed_ vs. Infected_mixed_, Uninfected_all_ vs. Uninfected_mixed_, Infected_all_ vs. Infected_mixed_) were performed with the *contrast* function from package *emmeans*^68^. P values were adjusted using the Sidak method for multiple comparisons.

GLMMs models were compared to reduced models using likelihood ratio tests to assess the significance of predictors using the function *drop1* in R. Assumptions of GLMMs were verified using model diagnostic tests and plots implemented in the package *DHARMa*.

## Supporting information

Mov S1. Nictating nematode attaches to an ant

Table S1. DE genes in the PPG

Table S2. No DE genes in the brain

Table S3. GO term enrichment analysis of the DE genes in PPG

## Authors’ contributions

Z.L.: sample collection, data curation, data analysis, writing—original draft, writing—review and editing; E.F.: data acquisition, data analysis (chemistry), writing—review and editing; F.A.: data acquisition, data analysis (micro-CT); V.B.: Data acquisition; T.OH.: data acquisition, data analysis (behaviour); D.J.P.: data analysis (sequencing), writing—review and editing; T.S.: data acquisition, data analysis (chemistry), writing—review and editing; E.E.: data acquisition, data analysis (micro-CT), writing—review and editing; Y.U.: conceptualization, funding acquisition, methodology, sample collection, project administration, resources, supervision, validation, writing—original draft, writing—review and editing. All authors contributed, reviewed and approved the manuscript for publication.

## Acknowledgements

The authors thank Giacomo Alciatore for assistance with automated tracking, Daniel Knebel, Alexandre Courtiol, and Grit Kunert for advice on data analysis, Hugo Darras and Jason Buser for sharing ant samples, OIST Imaging Section for access to the Micro-CT Scanner, Christine La Mendola for assistance with library preparation, Alexandra Bezler and the Gönczy lab for advice on nematode maintenance, and members of the Social Behaviour group for comments on the manuscript. Sequencing was performed at the Genomics Technologies Facility of the University of Lausanne.

## Data Availability

The behavioural and chemical data generated in this study, as well as the R scripts used for data analysis have been deposited in an Edmond repository (https://doi.org/10.17617/3.I6NKXM). The RNAseq data have been deposited at NCBI’s sequence read archive under the following BioProject accession: PRJNA791185. RNAseq accession numbers are in Tab. S6. Accession numbers for the Sanger sequences of nematodes isolates are: OP964514 (*P. longicornis*), OP964515 (*L. niger*), OP964516 (*O. biroi*).

## References

1. Graystock, P., Goulson, D. & Hughes, W. O. H. Parasites in bloom: flowers aid dispersal and transmission of pollinator parasites within and between bee species. Proc. R. Soc. B-Biol. Sci. 282, 20151371 (2015).

2. Albery, G. F., Becker, D. J., Kenyon, F., Nussey, D. H. & Pemberton, J. M. The Fine-Scale Landscape of Immunity and Parasitism in a Wild Ungulate Population. Integr. Comp. Biol. 59, 1165–1175 (2019).

3. de Bekker, C. et al. Species-specific ant brain manipulation by a specialized fungal parasite. BMC Evol. Biol. 14, 166 (2014).

4. Stroeymeyt, N. et al. Social network plasticity decreases disease transmission in a eusocial insect. Science 362, 941–945 (2018).

5. Stockmaier, S. et al. Infectious diseases and social distancing in nature. Science 371, eabc8881 (2021).

6. Hudson, P. J. The Ecology of Wildlife Diseases. (Oxford University Press, 2002).

7. Ezenwa, V. O. Habitat overlap and gastrointestinal parasitism in sympatric African bovids. Parasitology 126, 379–388 (2003).

8. Wood, M. J. et al. Within-population variation in prevalence and lineage distribution of avian malaria in blue tits, Cyanistes caeruleus. Mol. Ecol. 16, 3263–3273 (2007).

9. Plowright, R. K., Sokolow, S. H., Gorman, M. E., Daszak, P. & Foley, J. E. Causal inference in disease ecology: investigating ecological drivers of disease emergence. Front. Ecol. Environ. 6, 420–429 (2008).

10. Hawley, D. M. & Altizer, S. M. Disease ecology meets ecological immunology: understanding the links between organismal immunity and infection dynamics in natural populations. Funct. Ecol. 25, 48–60 (2011).

11. Wilson, E. O. The Ergonomics of Caste in the Social Insects. Am. Nat. 102, 41–66 (1968).

12. Mersch, D. P., Crespi, A. & Keller, L. Tracking Individuals Shows Spatial Fidelity Is a Key Regulator of Ant Social Organization. Science 340, 1090–1093 (2013).

13. Cremer, S., Armitage, S. A. O. & Schmid-Hempel, P. Social Immunity. Curr. Biol. 17, R693– R702 (2007).

14. Schmid-Hempel, P. Parasites in Social Insects. (Princeton University Press, 1998).

15. Blatrix, R. Task Allocation Depends on Matriline in the Ponerine Ant Gnamptogenys striatula Mayr. J. Insect Behav. 10 (2000).

16. Armitage, S. A. O. & Boomsma, J. J. The effects of age and social interactions on innate immunity in a leaf-cutting ant. J. Insect Physiol. 56, 780–787 (2010).

17. Bull, J. C. et al. A Strong Immune Response in Young Adult Honeybees Masks Their Increased Susceptibility to Infection Compared to Older Bees. Plos Pathog. 8, e1003083 (2012).

18. Invernizzi, C., Peñagaricano, F. & Tomasco, I. H. Intracolonial genetic variability in honeybee larval resistance to the chalkbrood and American foulbrood parasites. Insectes Sociaux 56, 233– 240 (2009).

19. Ravary, F. & Jaisson, P. The reproductive cycle of thelytokous colonies of Cerapachys biroi Forel (Formicidae, Cerapachyinae). Insectes Sociaux 49, 114–119 (2002).

20. Ulrich, Y., Saragosti, J., Tokita, C. K., Tarnita, C. E. & Kronauer, D. J. C. Fitness benefits and emergent division of labour at the onset of group living. Nature 560, 635–638 (2018).

21. Janet, C. Sur les nématodes des glandes pharyngiennes des fourmis (Pelodera sp.). Comptes Rendus Acad. Sci. 117, 700–703 (1893).

22. Wahab, A. Untersuchungen über Nematoden in den drüsen des kopfes der Ameisen (Formicidae). Z. Für Morphol. Ökol. Tiere 52, 33–92 (1962).

23. Nickle, W. & Ayre, G. Caenorhabditis dolichura (A. Schneider, 1866) Dougherty (Rhabditidae, Nematoda) in the head glands of the ants Camponotus herculeanus (L.) and Acanthomyops claviger (Roger) in Ontario. in Proceedings of the entomological society of ontario vol. 96 96 (1965).

24. Markin George. P. & McCoy, C. W. The Occurrence of a Nemotode, Diploscapter lycostoma, in the Pharyngeal Glands of the Argentine Ant, Iridomyrmex humilis. Ann. Entomol. Soc. Am. 61, 505–509 (1968).

25. Köhler, A. Nematodes in the heads of ants associated with sap flux and rotten wood. Nematology 14, 191–198 (2012).

26. Zhao, Z. Q. et al. Diploscapter formicidae sp. n. (Rhabditida: Diploscapteridae), from the ant Prolasius advenus (Hymenoptera: Formicidae) in New Zealand. Nematology 15, 109–123 (2013).

27. Richter, A. et al. The cephalic anatomy of workers of the ant species Wasmannia affinis (Formicidae, Hymenoptera, Insecta) and its evolutionary implications. Arthropod Struct. Dev. 49, 26–49 (2019).

28. Hölldobler, B. & Wilson, E. O. The Ants. (Harvard University Press, 1990).

29. Vinson, S. B., Philipps, S. A. & Williams, H. J. The Function of the Post-pharyngeal Glands of the Red Imported Fire Ant, Solenopsis invicta Buren. J. Insect Physiol. 26, 645–650 (1980).

30. Bagneres, A. G. & Morgan, E. D. The Postpharyngeal Glands and the Cuticle of Formicidae Contain the Same Characteristic Hydrocarbons. Experientia 47, 106–111 (1991).

31. Sprenger, P. P. & Menzel, F. Cuticular hydrocarbons in ants (Hymenoptera: Formicidae) and other insects: how and why they differ among individuals, colonies, and species. Myrmecol. News 30, 1–26 (2020).

32. Poinar, G. Nematode Parasites and Associates of Ants: Past and Present. Psyche J. Entomol. 2012, 1–13 (2012).

33. Campbell, J. F. & Gaugler, R. Nictation Behaviour and Its Ecological Implications in the Host Search Strategies of Entomopathogenic Nematodes (Heterorhabditidae and Steinernematidae). Behaviour 126, 155–169 (1993).

34. Lee, H. et al. Nictation, a dispersal behavior of the nematode Caenorhabditis elegans, is regulated by IL2 neurons. Nat. Neurosci. 15, 107–112 (2012).

35. Blaxter, M. L. et al. A molecular evolutionary framework for the phylum Nematoda. Nature 392, 71–75 (1998).

36. Peregrine, D. J., Mudd, A. & Cherrett, J. M. Anatomy and preliminary chemical analysis of the post-pharyngeal glands of the leaf-cutting ant, Acromyrmex octospinosus (Reich.) (Hym., Formicidae). Insectes Sociaux 20, 355–363 (1973).

37. Matyash, V. et al. Sterol-Derived Hormone(s) Controls Entry into Diapause in Caenorhabditis elegans by Consecutive Activation of DAF-12 and DAF-16. PLOS Biol. 2, e280 (2004).

38. Ulrich, Y. et al. Response thresholds alone cannot explain empirical patterns of division of labor in social insects. PLOS Biol. 19, e3001269 (2021).

39. Chandra, V., Gal, A. & Kronauer, D. J. C. Colony expansions underlie the evolution of army ant mass raiding. Proc. Natl. Acad. Sci. 118, (2021).

40. Leclerc, J.-B. & Detrain, C. Loss of attraction for social cues leads to fungal-infected Myrmica rubra ants withdrawing from the nest. Anim. Behav. 129, 133–141 (2017).

41. Trinh, T., Ouellette, R. & de Bekker, C. Getting lost: the fungal hijacking of ant foraging behaviour in space and time. Anim. Behav. 181, 165–184 (2021).

42. Heinze, J. & Walter, B. Moribund Ants Leave Their Nests to Die in Social Isolation. Curr. Biol. 20, 249–252 (2010).

43. Naug, D. & Camazine, S. The Role of Colony Organization on Pathogen Transmission in Social Insects. J. Theor. Biol. 215, 427–439 (2002).

44. Stroeymeyt, N., Casillas-Pérez, B. & Cremer, S. Organisational immunity in social insects. Curr. Opin. Insect Sci. 5, 1–15 (2014).

45. Ulrich, Y. & Schmid-Hempel, P. The distribution of parasite strains among hosts affects disease spread in a social insect. Infect. Genet. Evol. J. Mol. Epidemiol. Evol. Genet. Infect. Dis. 32, 348– 353 (2015).

46. Ravary, F. & Jaisson, P. Absence of individual sterility in thelytokous colonies of the ant Cerapachys biroi Forel (Formicidae, Cerapachyinae). Insectes Sociaux 51, 67–73 (2004).

47. Jud, S. L., Knebel, D. & Ulrich, Y. Intergenerational genotypic interactions drive collective behavioural cycles in a social insect. Proc. R. Soc. B Biol. Sci. 289, 20221273 (2022).

48. Kronauer, D. J. C., Pierce, N. E. & Keller, L. Asexual reproduction in introduced and native populations of the ant Cerapachys biroi. Mol. Ecol. 21, 5221–5235 (2012).

49. Stiernagle, T. Maintenance of C. elegans. WormBook 1–11 (2006).

50. Holterman, M. et al. Phylum-Wide Analysis of SSU rDNA Reveals Deep Phylogenetic Relationships among Nematodes and Accelerated Evolution toward Crown Clades. Mol. Biol. Evol. 23, 1792–1800 (2006).

51. Löytynoja, A. & Goldman, N. An algorithm for progressive multiple alignment of sequences with insertions. Proc. Natl. Acad. Sci. U. S. A. 102, 10557–10562 (2005).

52. Talavera, G. & Castresana, J. Improvement of phylogenies after removing divergent and ambiguously aligned blocks from protein sequence alignments. Syst. Biol. 56, 564–577 (2007).

53. Stamatakis, A. RAxML version 8: a tool for phylogenetic analysis and post-analysis of large phylogenies. Bioinformatics 30, 1312–1313 (2014).

54. Libbrecht, R., Oxley, P. R., Keller, L. & Kronauer, D. J. C. Robust DNA Methylation in the Clonal Raider Ant Brain. Curr. Biol. 26, 391–395 (2016).

55. Bolger, A. M., Lohse, M. & Usadel, B. Trimmomatic: a flexible trimmer for Illumina sequence data. Bioinformatics 30, 2114–2120 (2014).

56. McKenzie, S. K. & Kronauer, D. J. C. The genomic architecture and molecular evolution of ant odorant receptors. Genome Res. 28, 1757–1765 (2018).

57. Bray, N. L., Pimentel, H., Melsted, P. & Pachter, L. Near-optimal probabilistic RNA-seq quantification. Nat. Biotechnol. 34, 525–527 (2016).

58. Soneson, C., Love, M. I. & Robinson, M. D. Differential analyses for RNA-seq: transcript-level estimates improve gene-level inferences. F1000Research 4, 1521 (2015).

59. Robinson, M. D., McCarthy, D. J. & Smyth, G. K. edgeR: a Bioconductor package for differential expression analysis of digital gene expression data. Bioinformatics 26, 139–140 (2010).

60. Benjamini, Y. & Hochberg, Y. Controlling the False Discovery Rate: A Practical and Powerful Approach to Multiple Testing. J. R. Stat. Soc. Ser. B Methodol. 57, 289–300 (1995).

61. Götz, S. et al. High-throughput functional annotation and data mining with the Blast2GO suite. Nucleic Acids Res. 36, 3420–3435 (2008).

62. Alexa, A., Rahnenführer, J. & Lengauer, T. Improved scoring of functional groups from gene expression data by decorrelating GO graph structure. Bioinformatics 22, 1600–1607 (2006).

63. Supek, F., Bošnjak, M., Škunca, N. & Šmuc, T. REVIGO Summarizes and Visualizes Long Lists of Gene Ontology Terms. PLOS ONE 6, e21800 (2011).

64. Gal, A., Saragosti, J. & Kronauer, D. J. anTraX, a software package for high-throughput video tracking of color-tagged insects. eLife 9, e58145 (2020).

65. R Core Team. R: A Language and Environment for Statistical Computing. (R Foundation for Statistical Computing, 2020).

66. Therneau, T. M. coxme: Mixed Effects Cox Models. (2022).

67. Oksanen, J. et al. vegan: Community Ecology Package. (2022).

68. Lenth, R. V. emmeans: Estimated Marginal Means, aka Least-Squares Means. (2022).

69. Brooks, M. E. et al. glmmTMB Balances Speed and Flexibility Among Packages for Zero-inflated Generalized Linear Mixed Modeling. R J. 9, 378–400 (2017).

