## Supplementary Materials for "Behavioural individuality determines infection risk in clonal ant colonies"

Authors: Li, Z.^1,2^, Frank, E.T.^3^, Oliveira-Honorato, T.^4^, Azuma, F^5^., Bachmann, V^2^., Parker, D. J.^6^, Schmitt, T.^3^, Economo, E.^5^, Ulrich, Y.^1,2,4^

^1^Max Planck Institute for Chemical Ecology, Jena, Germany

^2^Institute of Integrative Biology, ETH Zürich, Zürich, Switzerland

^3^Department of Animal Ecology and Tropical Biology, Biocentre, University of Würzburg, Würzburg, Germany

^4^Department of Ecology and Evolution, University of Lausanne, Lausanne, Switzerland

^5^Biodiversity and Biocomplexity Unit, Okinawa Institute of Science and Technology Graduate University, Onna, Japan

^6^School of Natural Sciences, Bangor University, Bangor, United Kingdom

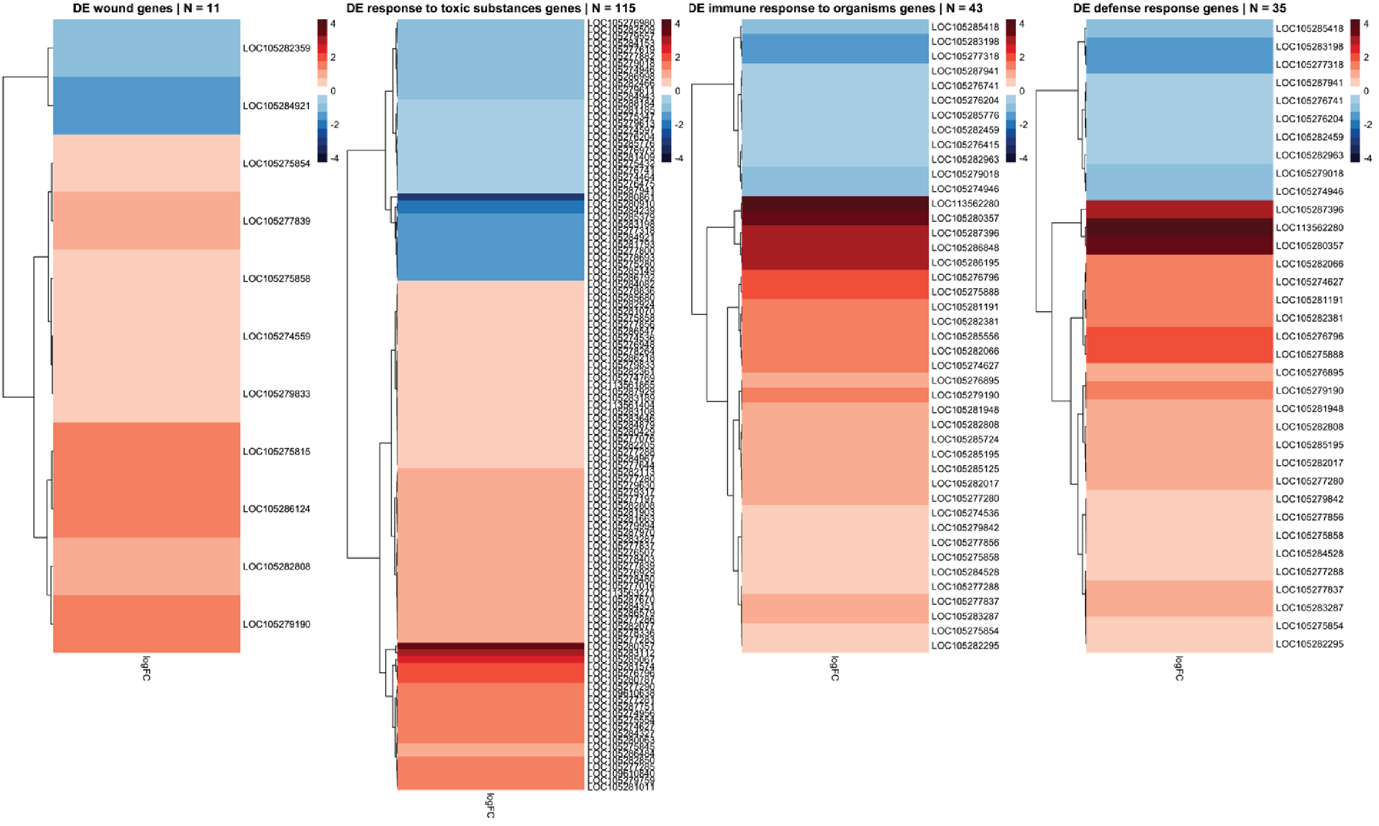

**Fig S1.** Differential expression of immune gene clusters between non-infected and infected PPGs. Red: higher expression in infected samples, blue: lower expression.

**Mov S1.** Nictating nematode attaches to an ant

See *MovS1_Nictating_nematode_attaches_to_an_ant.mp4*

### Tables

**Table S1.** DE genes in the PPG

See *TableS1_ PPG_DE_with_immune_beh_interestother_df_nrDroso.csv*

**Table S2.** No DE genes in the brain

See *TableS2_fit_BRN_F2_inf_DE_out.csv*

**Table S3.** GO term enrichment analysis of the DE genes in PPG

See *TableS3_Revigo_PPG_BP_005_Small_a.xlsx*

**Table S4.** Median relative abundance ± SD of all CHCs in ants with different genotypes (B, L) and infection status (uninfected, infected).

| **Compound** | **B_uninfected_** (n=10) | **B_infected_** (n=10) | **L_uninfected_** (n=10) | **L_infected_** (n=10) |
| --- | --- | --- | --- | --- |
| C25 | 1.0±0.2 | 0.4±0.2 | 1.0±0.2 | 0.0±0.0 |
| 11-MeC25/13-MeC25 | 3.3±0.4 | 2.0±0.5 | 1.6±0.2 | 2.9±0.8 |
| 5-MeC25 | 1.9±0.5 | 1.3±0.5 | 0.6±0.1 | 1.4±0.6 |
| 2-MeC25 | 7.0±0.4 | 5.5±0.4 | 8.4±0.2 | 4.5±0.5 |
| C26 | 0.4±0.6 | 0.3±0.3 | 0.6±0.1 | 0.0±0.0 |
| 2-MeC26 | 5.3±0.2 | 6.5±0.3 | 3.1±0.2 | 5.7±0.4 |
| C27 | 14.2±1.2 | 9.7±1.1 | 15.3±1.8 | 7.5±2.6 |
| 13-MeC27 | 13.1±1.1 | 12.0±0.2 | 13.9±0.1 | 11.5±0.5 |
| 11,15-diMeC27 | 51.8±3.2 | 59.6±3.2 | 54.8±1.5 | 64.0±3.4 |
| 3-MeC27 | 0.0±0.0 | 0.0±0.0 | 0.6±0.2 | 0.7±0.5 |

**Table S5.** X-ray micro-CT scan parameters.

| **Species** | **Specimen identifier** | **Voxel size** | **Exposure time (s)** | **Projections** | **Voltage (kV)** | **Power (W)** | **Source distance (mm)** | **Detector distance (mm)** | **Magnification** |
| --- | --- | --- | --- | --- | --- | --- | --- | --- | --- |
| *Lasius niger* | CASENT0744001 | 1.164 | 22 | 1601 | 40 | 3 | -12.504 | 60.002 | 4 |
| *Ooceraea biroi* | CASENT0741215 | 0.995 | 13 | 3201 | 40 | 3 | -9.516 | 55.019 | 4 |
| *Paratrechina longicornis* | CASENT0744371 | 0.778 | 24.5 | 1601 | 40 | 3 | -10.043 | 77.037 | 4 |

**Table S6.** Read counts and accession numbers for sequenced samples.

| **Sample Name** | **Tissue** | **Infection status** | **Raw reads** | **Trimmed reads** | **Mapped reads** | **SRA accession** |
| --- | --- | --- | --- | --- | --- | --- |
| Obir_BRN_I_rep2 | Brain | Infected | 20289459 | 19398729 | 13354218 | SRR17285162 |
| Obir_BRN_I_rep4 | Brain | Infected | 18353702 | 17586526 | 12783548 | SRR17285154 |
| Obir_BRN_I_rep6 | Brain | Infected | 17616117 | 16902695 | 12235276 | SRR17285148 |
| Obir_BRN_I_rep8 | Brain | Infected | 24303035 | 22959320 | 15826010 | SRR17285160 |
| Obir_BRN_NI_rep1 | Brain | Uninfected | 22676153 | 21646247 | 15233311 | SRR17285163 |
| Obir_BRN_NI_rep3 | Brain | Uninfected | 20565461 | 19756221 | 14353756 | SRR17285155 |
| Obir_BRN_NI_rep5 | Brain | Uninfected | 24824020 | 23803884 | 17407569 | SRR17285149 |
| Obir_BRN_NI_rep7 | Brain | Uninfected | 24411277 | 23512768 | 17719564 | SRR17285161 |
| Obir_PPG_I_rep2 | PPG | Infected | 18715309 | 17973940 | 12478900 | SRR17285150 |
| Obir_PPG_I_rep4 | PPG | Infected | 18754300 | 18090812 | 12187685 | SRR17285153 |
| Obir_PPG_I_rep6 | PPG | Infected | 20491749 | 19149551 | 13668256 | SRR17285158 |
| Obir_PPG_I_rep8 | PPG | Infected | 18310898 | 17618160 | 13177067 | SRR17285156 |
| Obir_PPG_NI_rep1 | PPG | Uninfected | 19101715 | 18383613 | 14485763 | SRR17285151 |
| Obir_PPG_NI_rep3 | PPG | Uninfected | 16605032 | 15979022 | 12686796 | SRR17285152 |
| Obir_PPG_NI_rep5 | PPG | Uninfected | 17770008 | 16957202 | 13142744 | SRR17285159 |
| Obir_PPG_NI_rep7 | PPG | Uninfected | 20854374 | 19986375 | 15554046 | SRR17285157 |
